# Functional gene embeddings improve rare variant polygenic risk scores

**DOI:** 10.1101/2024.07.22.604535

**Authors:** Shubhankar Londhe, Jonas Lindner, Zhifen Chen, Eva Holtkamp, Florian R. Hölzlwimmer, Francesco Paolo Casale, Felix Brechtmann, Julien Gagneur

## Abstract

Rare variant association testing is a powerful strategy for identifying effector genes underlying common traits. However, its effectiveness is limited by the scarcity of high-impact rare allele carriers, posing challenges for sensitivity and robustness. Here, we introduce FuncRVP, a rare variant association framework addressing this issue by leveraging functional information across genes. FuncRVP models the effects of rare variants as a weighted sum of gene impairment scores, with weights regularized through a prior based on functional gene embeddings. Modeling 41 quantitative traits from unrelated UK Biobank participants showed that FuncRVP consistently outperformed linear regressions on significantly associated genes and did so more effectively for traits with higher burden heritability. The framework demonstrated versatility, yielding consistent improvements across diverse gene embeddings. Moreover, FuncRVP generated more robust gene effect estimates and yielded more gene discoveries, especially among genetically constrained genes. These findings demonstrate the value of integrating functional information in rare variant association studies and showcase FuncRVP as a promising tool for enhancing phenotype prediction and gene discovery.

## Main

Population-scale whole-exome and whole-genome sequencing vastly expand the potential to unravel the genetic underpinnings of human traits and diseases. Such large-scale sequencing studies have unveiled that an average individual carries dozens of potentially deleterious rare germline variants^1^. Rare coding variants (allele frequency < 0.1%), on average, explain substantially less phenotypic variance than common variants and contribute only modestly to missing heritability^2^. However, they can exhibit large effects, aiding the discovery of effector genes with major roles in the molecular mechanisms underlying diseases and, therefore, prioritizing potential drug targets. For instance, the discovery of protective loss-of-function (LoF) variants in proprotein convertase subtilisin kexin 9 (*PCSK9*) led to the development of inhibitors for coronary heart disease^3–5^. Furthermore, rare variant-derived polygenic risk scores (PRS) not only allow identifying individuals at high disease risk but also better generalize across populations with different ethnicities, benefitting from a milder dependence on linkage disequilibrium (LD) structure compared to common variants^6^.

While genome-wide association studies have been instrumental in identifying common genetic variants associated with complex traits, they face challenges when analyzing rare variants due to the scarcity of allele carriers and increased multiple testing burden. To address this limitation, rare variant association tests (RVATs), including burden tests and variance-component tests such as the sequence kernel association test, aggregate rare variants within a gene or a genomic region^7–12^. To further improve rare variant association testing by leveraging rich and heterogenous variant annotations, we have recently developed DeepRVAT, a deep learning model that learns from genetic and phenotypic data how to integrate annotations of all rare variants into a trait-agnostic and individual-specific gene impairment score^13^. DeepRVAT models the phenotype as a weighted sum of gene impairment scores of significantly associated genes. Despite this simplifying assumption, DeepRVAT outperformed previous rare variant association testing approaches for gene discovery and phenotype prediction^13^.

Rare variant association testing studies often reveal functionally related genes associated with the same traits (see, for example ref.^6,14^). Leveraging this observation, gene-set RVATs perform burden tests or variance-component tests on a-priori-defined groups of genes, such as pathways^15–18^. Gene-set RVAT can show increased sensitivity in capturing pathways associated with traits^19^; however, this approach neither allows pinpointing individual gene associations nor deriving phenotype predictors.

Recently, functional gene embeddings, numerical vectors capturing gene functions, have been proposed as an alternative to gene sets to represent gene functions^20–22^. The embedding space is such that genes with similar functions are close to each other^20,23^. Since they are vector representations rather than sets, functional gene embeddings can retain some quantitative information about functional similarities and are more straightforwardly integrated into machine learning algorithms. Recently, the development of Polygenic Priority Score (PoPS) has shown the utility of functional gene embeddings to prioritize genes at associated loci to be helpful for genome-wide association studies^23^. However, their utility for rare variant association testing has not yet been explored.

Here, we propose to leverage functional gene embeddings to increase the power of RVAT while maintaining the capability of developing phenotype predictors and individual gene discovery. To this end, we developed a genotype-to-phenotype model we named FuncRVP (Functional Rare Variant Polygenic Risk Score). FuncRVP predicts phenotypes as a weighted sum of common- variant PRS and gene impairment scores for all genes, with gene weights regularized by a trait- specific prior that depends on a functional gene embedding. In the following, we present the model architecture, explore the modeling assumptions using 41 quantitative traits across 372,392 unrelated individuals of the UK Biobank, and assess the model robustness on held-out individuals. We demonstrate superior phenotype prediction against DeepRVAT-based predictors and improved robustness of gene effect estimates. We show that these improvements are robust to the choice of gene impairment scores and functional gene embeddings. Moreover, we assess the biological relevance of FuncRVP novel associations by correlating them with GWAS signals for the same traits and causal genes for rare Mendelian disorders with related phenotypes.

## Results

### The FuncRVP prior regularizes gene effects using functional gene embeddings

FuncRVP is a Bayesian model that predicts quantitative phenotypes from rare and common variants, utilizing a functional gene embedding to borrow information across genes (Fig. 1). In contrast to existing rare-variant-based polygenic risk scores^6,24^ FuncRVP does not select predictive genes using statistical testing. Instead, FuncRVP stands out by modeling the trait considering rare variant gene impairment from all genes jointly. Technically, this is robustly achieved by predicting phenotypes as a weighted sum of common-variant PRS and gene impairment scores for all genes, with gene weights regularized by a trait-specific prior that depends nonlinearly on a functional gene embedding. We made Gaussian assumptions to model the prior and the phenotype leading to a closed-form solution of the posterior of the phenotype. The model was implemented in the deep learning framework PyTorch and fitted by maximizing the posterior of the phenotype. Hyperparameter selection for the neural network used to model the prior and model evaluation was achieved through a train-validation-test split including 65%, 10%, and 25% of the individuals respectively (Methods). We used the validation split to select the best hyperparameters and parameters of the neural network. Once the neural network was selected and trained, we computed the gene effect posteriors using the union of the training and validation samples. These gene effect posteriors were then fixed and used to obtain phenotype predictions on the test split.

**Figure 1.**
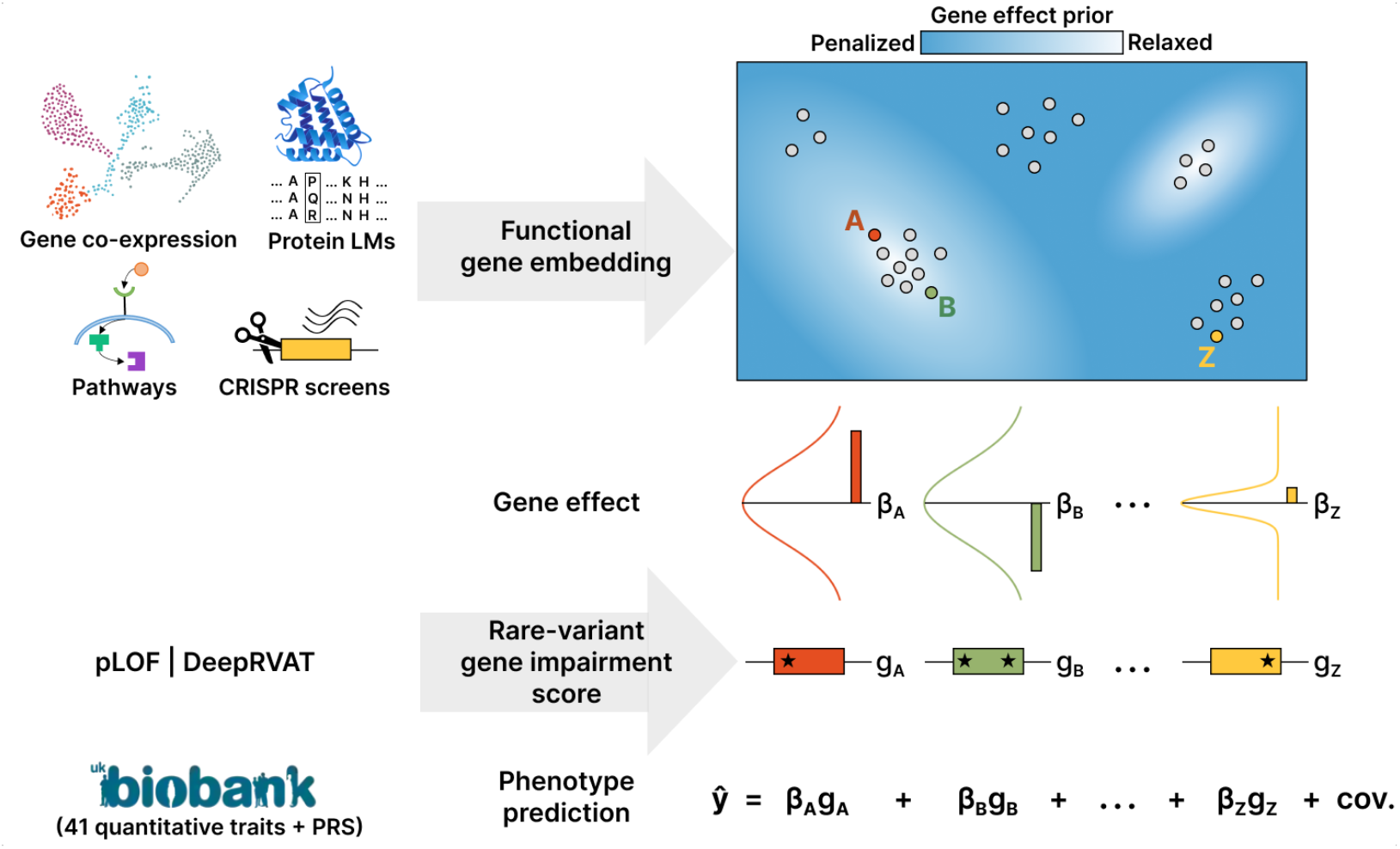
FuncRVP model overview. FuncRVP is a Bayesian model that predicts phenotype from rare and common variants. An individual’s phenotype is modeled as a weighted sum of their rare variant gene impairment scores and covariates, including precomputed common-variant polygenic risk scores (bottom). A prior, learned from data, constrains the magnitude of gene effects depending on their location in a functional gene embedding, resulting in relaxed regularizations on areas densely populated with trait-associated genes (e.g., gene A and gene B). We investigated several functional gene embeddings based on various functional genomics data and rare-variant impairment scores based on putative loss of function variants (pLOF)^25^ and on the integrative gene impairment score DeepRVAT^24^.

We trained FuncRVP models for 41 quantitative phenotypes (Table S1) on 372,392 unrelated individuals with European ancestry in the UK Biobank who underwent whole-exome sequencing^26^. As gene impairment scores, we initially used the DeepRVAT scores precomputed on the UK Biobank^13^. We considered 24 covariates including polynomial terms of age and sex, genetic principal components, and a common-variant polygenic risk score (Methods, Table S2). To avoid data leakage, DeepRVAT uses a cross-validation scheme that ensures that for every individual, the gene impairment scores were derived from models that were not trained using that individual. Moreover, we initially considered as functional gene embeddings the concatenation of two best-performing gene embeddings investigated by Brechtmann and colleagues^20^. One of these two embeddings was the “Omics” embedding, which is based on gene expression data, genome-wide CRISPR screen data, and a protein language model. The other embedding was a low-dimensional representation of the gene feature matrix from the PoPS study^23^, which is based on gene expression and pathway annotations and which retained the same performance as the PoPS original feature matrix across a range of benchmarks^20^.

We assessed two alternative structures for the prior: one in which the expectation of the weights depended on the embedding and another one in which the variance of the weights depended on the embedding. Across all traits, we found that modeling the variance consistently led to substantially more accurate and robust phenotype predictors on held-out data (Fig. 2A). These results indicate that the position in the embeddings was predictive of whether genes affect a phenotype but not of the value of the effect. Another possibility is that the model lacked sufficient data to reliably capture directional information, though it could still identify regions in the embedding space enriched with genes exhibiting large effects. One example illustrating this phenomenon is shown in Fig. 2B, which shows the distribution of HDL cholesterol levels among individuals with a putative loss of function variant for the Cholesteryl ester transfer protein *CETP* and its 24 nearest genes in the functional embedding. Like *CETP*, 4 out of the 24 nearest neighbors have been reported to be among the 42 significantly associated genes with HDL cholesterol levels^27^ (Enrichment 94-fold, *P* < 2.7 × 10^−7^, Fisher test). However, the effects of these genes vary widely in amplitude and direction.

**Figure 2.**
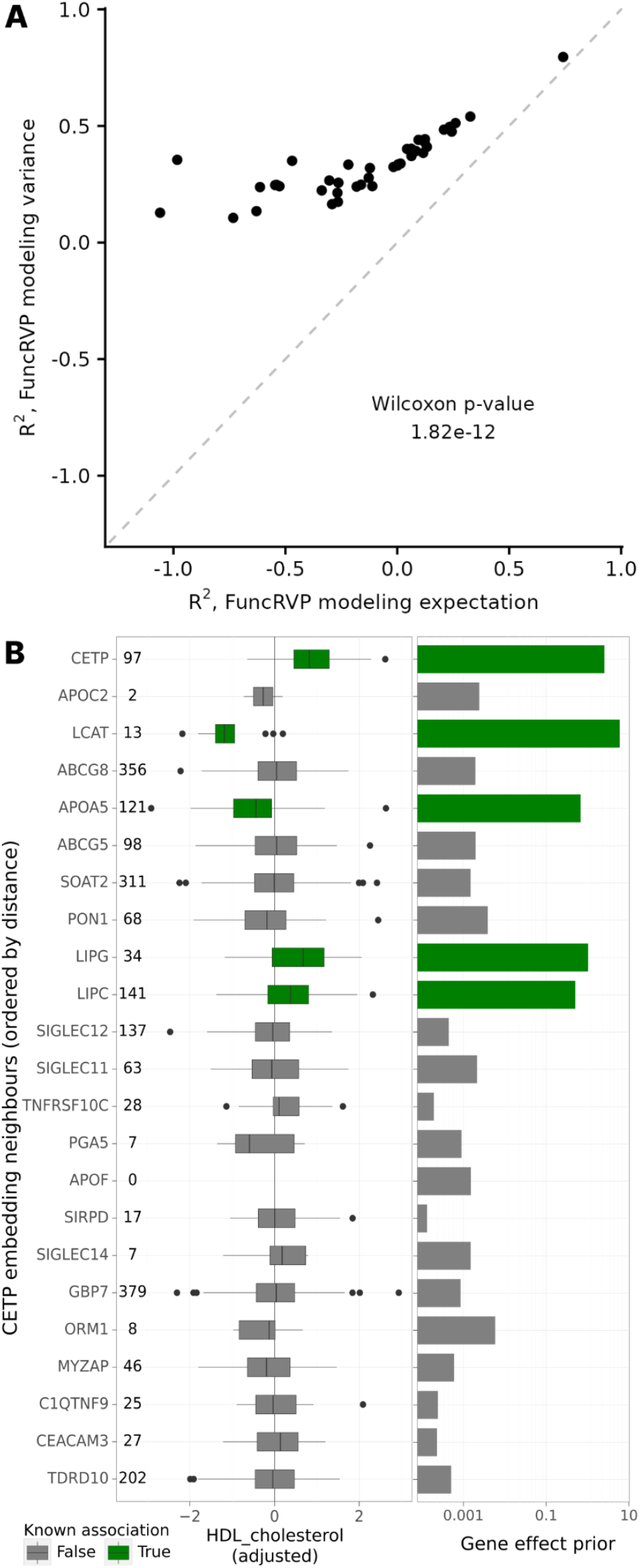
Assessing FuncRVP model assumptions. **A)** Explained variance on held-out data using a prior on the gene effect variance (y-axis) against a prior on gene effect expectations (x-axis) across 41 traits. (*P* < 2×10^−12^, paired Wilcoxon test). **B)** The gene *CETP* and its nearest neighbors in the embedding space are ordered by increasing distance. Left. Distribution of HDL cholesterol values (adjusted for common variants and covariates) among individuals with a putative loss-of-function (pLOF) variant in a given gene. Center line, median; box limits, first and third quartiles; whiskers span all data within 1.5 interquartile ranges of the lower and upper quartiles. Right. The variance of the prior learned by the model (y-axis) is plotted for each of these genes for the trait HDL cholesterol. Data for genes reported to be significantly associated by previous RVAT studies^27,28^ are in green and in gray otherwise.

### FuncRVP improves phenotype prediction

Having established the model’s prior structure, we next assessed the added value of FuncRVP for phenotype prediction on held-out test samples. As a state-of-the-art comparison point, we used linear models on DeepRVAT gene impairment scores of significantly associated genes, as described earlier^13^. Since the largest fraction of phenotypic variance is typically explained by common variants, we used as a comparison metric the relative ΔR^2^ compared to a common-variant baseline model linearly integrating polygenic risk scores, age, and sex covariates (Methods). We observed significant improvements in phenotype prediction for 25 out of 41 traits, with no single trait showing a significantly worse performance for FuncRVP (*P* < 0.05; Methods; Fig. 3A). The traits for which FuncRVP improved showed greater burden heritability, which is an estimated fraction of phenotypic variance explained by minor allele burden^2^, a larger number of genes previously found to be associated by RVATs^27,28^, and larger number of significant GWAS loci (Methods, Fig. 3B-D). Collectively, these results show that FuncRVP provides improved phenotypic predictors. The model best improves for traits for which the genetic basis is stronger, presumably because this helps learn the prior function.

**Figure 3.**
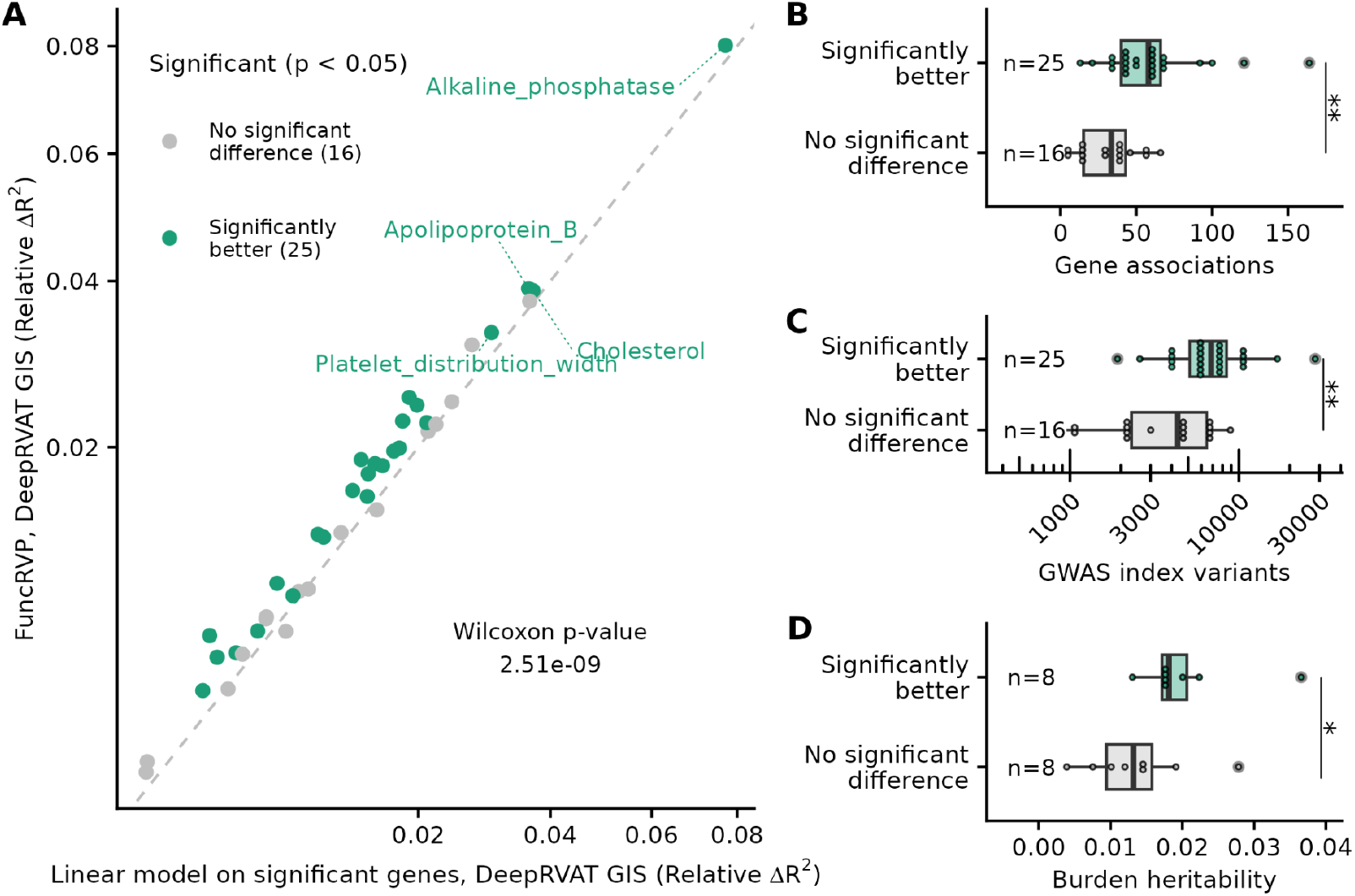
FuncRVP improves phenotype prediction, particularly for phenotypes with a stronger genetic basis. **A)** Relative improvement in phenotype prediction of FuncRVP (y-axis) compared to a linear regression on gene impairment scores of significant genes (x-axis) for 41 phenotypes. The relative fit improvement was computed as the improvement in R^2^ relative to a baseline linear model using only the PRS and covariates (Methods). Trait-wise significance was determined by case bootstrapping (Methods). Overall significance was determined through a paired Wilcoxon test. **B)** Burden heritability scores for 16 traits^2^, compared between traits that performed significantly better with FuncRVP and those for which FuncRVP showed no significant improvement. **C)** Comparison of the number of gene-trait associations found for each trait by previous RVASs^27,28^ across three categories: traits where FuncRVP performed significantly better, traits where there was no significant difference, and traits where FuncRVP performed significantly worse. **D)** As in C using the number of GWAS index variants associated with each trait across the three categories. For B-D) Center line, median; box limits, first and third quartiles; whiskers span all data within 1.5 interquartile ranges of the lower and upper quartiles. Significance was determined with the one-sided Wilcoxon test.

### FuncRVP is robust to functional gene embedding and gene impairment score choices

We next investigated alternative functional gene embedding choices. To cover functional information captured by protein sequence, we considered the protein language model ESM2^29^. To cover functional information reflected by gene regulation, we considered gene2vec, an embedding derived from gene expression data only^21^. Furthermore, we built a gene embedding using the sequence representation of gene transcription start sites of the Enformer^30^, a sequence- based predictor of gene expression and chromatin states for tens of cell lines and culture conditions (Method). For comparability, all these embeddings were reduced to a common dimension by projecting on their respective 512 first principal components, except gene2vec, because it originally had only 200 dimensions. Regardless of the embedding choice, FuncRVP improved phenotype prediction for more traits than it deteriorated, compared to linear models based on burden-test significant genes (Fig. 4). The strongest improvement was observed for embeddings capturing gene regulation in some way: Omics+PoPS, gene2vec, and Enformer all showed significant improvements on at least 25 out of the 41 traits. In comparison, the embedding based on protein sequence only, ESM2, significantly improved only for 19 traits. Importantly, all those models significantly improved over the linear model based on burden-test significant genes (paired Wilcoxon test, *P* < 0.05). In contrast, a FuncRVP model based on a randomly generated embedding of 512 dimensions was significantly worse than the linear model based on burden- test significant genes (Fig. S1), demonstrating that the FuncRVP improvements were not driven by a mere regularization but did require and leverage functional gene information.

**Figure 4.**
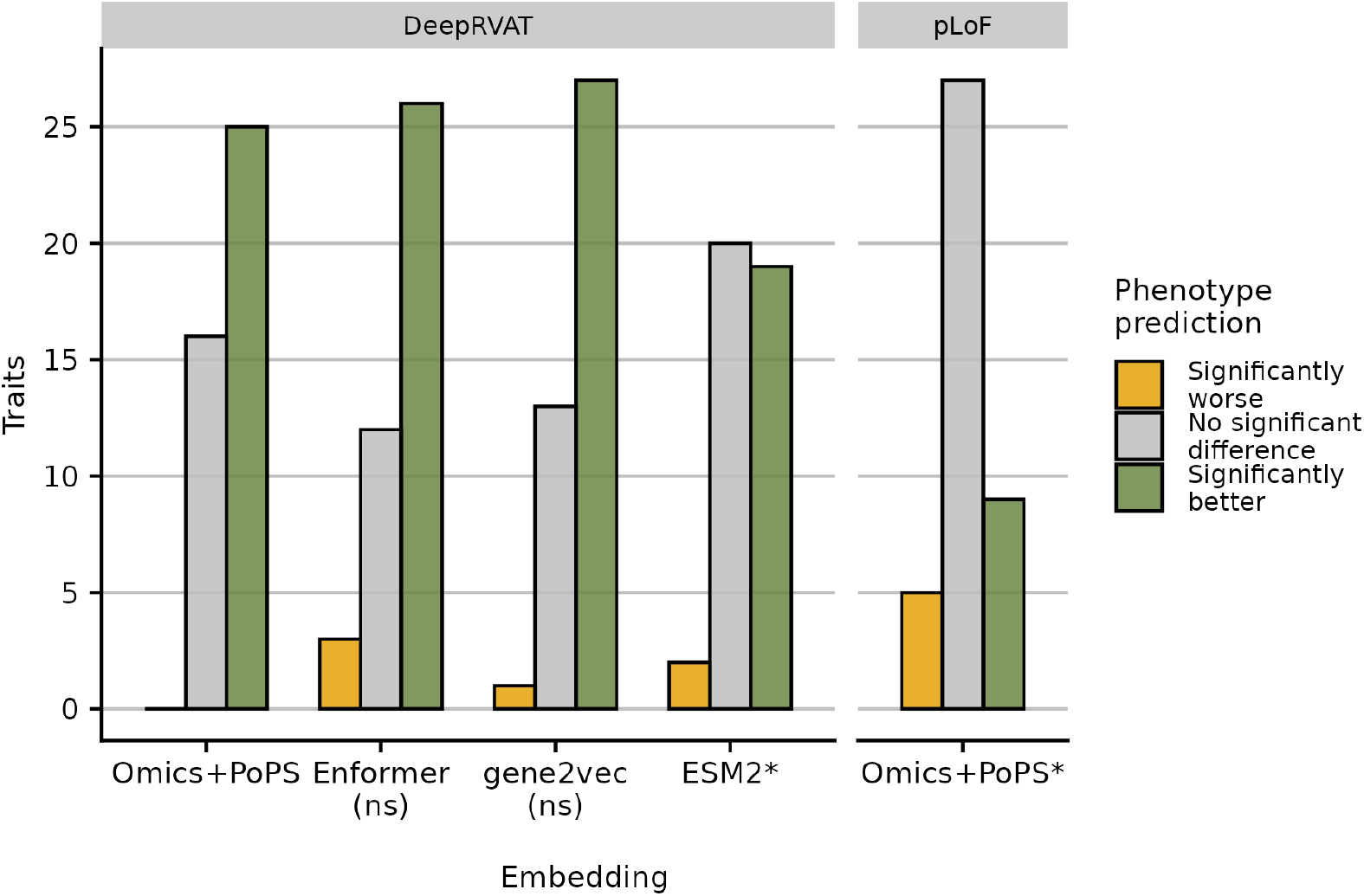
Robustness to choice of gene embedding and gene impairment scores. Number of traits with significant phenotype prediction improvement (green), not significant (grey), and significantly worse (orange) when using FuncRVP compared to a linear model on burden-test significant genes. Trait-wise significance was determined by case bootstrapping as in Fig. 3A. Data is shown for various combinations of functional gene embeddings (x-axis) and gene impairment scores (facets). For all combinations, phenotype predictions using FuncRVP were more often improved than worsened. Across the combinations, the Omics+PoPS gene embedding combined with the DeepRVAT scores performed similarly as the gene2vec or the Enformer embeddings combined with the DeepRVAT scores and significantly better than any other combinations (two-sided paired Wilcoxon test between trait-level relative ΔR^2^, akin to Fig. 3A, *P* < 0.05, (ns) = not significantly different, * = significantly worse).

Altogether, these results show that functional gene embeddings can be leveraged to improve the integration of rare variants into phenotype predictors. To this end, expression similarity appears more informative than protein sequence similarity.

Moreover, we found that FuncRVP improved on 9 traits and was worse on 5 compared to linear regression on burden-test significant genes when using the total number of putative loss-of- function (pLOF) variants instead of DeepRVAT for gene impairment scoring (Fig. 4). This indicates that FuncRVP is robust to the choice of gene impairment score.

### Improved estimation of gene effects using FuncRVP

Having shown that FuncRVP improves phenotype predictions, we next asked whether it helped provide robust estimates of gene effects on phenotypes compared to linear regression on burden-test significant genes as implemented by DeepRVAT. Comparing these methods is inherently challenging due to their different statistical paradigms: Bayesian for FuncRVP and frequentist for burden testing. To not bias the benchmark toward FuncRVP, we defined benchmark gene effect values using standard, non-regularized linear regression applied to the held-out test set samples (Fig. 5A).

**Figure 5.**
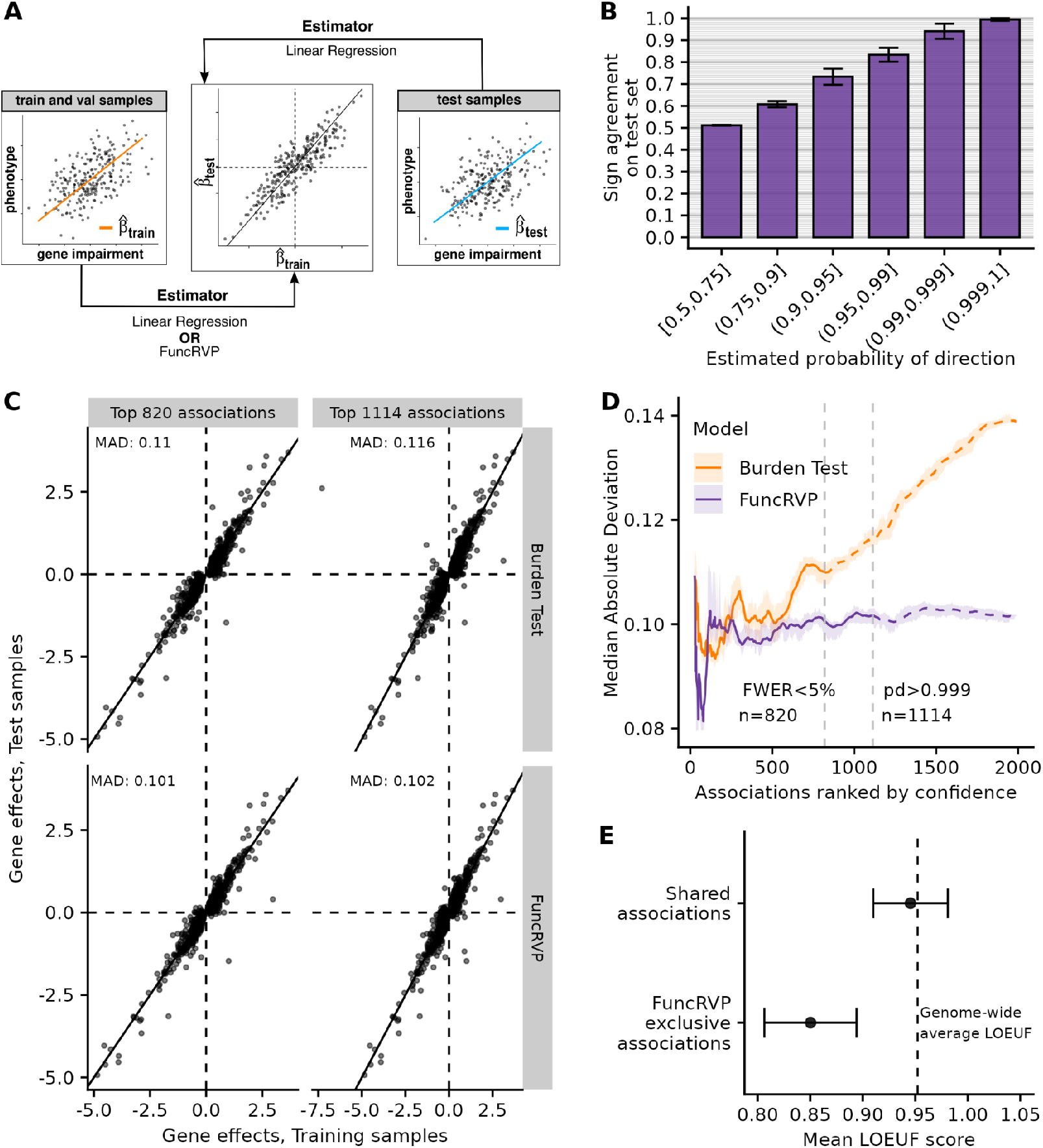
FuncRVP is more sensitive and yields more robust gene effect estimates than burden testing. **A)** Strategy for assessing the accuracy of gene effect estimates using held-out test samples. Gene effect estimates from FuncRVP or linear regression are compared against those obtained on held-out test samples using linear regression. **B)** Distribution across 41 traits of the proportion of sign agreements between FuncRVP gene effects estimated on the training phase samples and gene effects estimated by linear regression on the test samples. Error bars denote 95% confidence intervals of the mean across all 41 traits. **C)** Gene effect estimates on the test samples (y-axis) against training phase samples (x-axis). Associations up to the discovery thresholds of both methods were compared (820 trait-wise Bonferroni significant associations, *P* < 0.05, found by linear regression, and 1,114 associations with probability of direction > 0.999 found by FuncRVP). The median absolute deviation is indicated for each scatter plot. **D)** Median absolute deviation of predicted gene effects (y-axis) up to a given rank (x-axis) across all 41 traits. Genes are ranked by p-value for the burden test and probability of direction for FuncRVP. The ribbon represents a 95% equi-tailed confidence interval estimated by the jackknife across traits. **E)** The mean LOEUF score of genes (x-axis) in exclusive associations of FuncRVP and shared associations (y-axis). An association is non-exclusive if it is captured by both FuncRVP and a burden test and exclusive if found only by FuncRVP. Error bars denote 95% confidence intervals of the mean across all associations in the category (*P* < 10^−3^, two-sided t-test). The dashed vertical line denotes the average LOEUF score computed across all protein-coding genes.

Burden-test associations were ranked by increasing p-values, whereas FuncRVP associations were ranked by decreasing probability of direction, a Bayesian metric that quantifies the certainty about the sign of an effect (reviewed in ref.^31^).

Firstly, we assessed the agreement in effect direction estimated by FuncRVP using the training phase samples, i.e. the union of the train and validation splits, compared to those obtained by linear regression in the test samples. Generally, the probability of direction overestimated the empirical sign agreements. This overestimation could be either due to model miscalibration, or noise in the empirical effect estimations in the test samples, or a combination of both. Nonetheless, the agreement in direction increased as the probability of direction increased, showing that ranking based on the probability of direction is informative about the robustness of the estimates. Moreover, the directions of effects estimated by FuncRVP with a probability of direction larger than 0.999 perfectly agreed with those in the test samples for almost all traits (Fig. 5B). Accordingly, we considered a probability of direction of 0.999 as the minimum threshold for a FuncRVP association to be identified as discovered. Across the 41 traits, this cutoff yielded 1,114 associations, i.e. on average, 27.2 associations per trait (+/- 21.0 standard deviation). In comparison, the DeepRVAT-based burden test led to 820 significant associations using the training phase samples (Family-wise error rate < 0.05, trait-wise Bonferroni correction).

Next, we quantitatively assessed the agreement between the gene effects estimated in the training phase samples and the benchmark gene effect values from the test split. On median, the absolute differences between the estimates and the benchmark values were smaller for FuncRVP than for linear regression, both among the top 820 associations and among the top 1,114 associations (Fig. 5C). Moreover, the improvement was significant independently of association rank cutoff (Fig. 5D). Altogether, these results demonstrate that FuncRVP improves effect estimations over burden-test based approaches.

Genes with low tolerance to loss function variants^1^ are an interesting class of genes because they are more likely to have functional importance. However, as they are, by definition, depleted for high-impact variants, these genes particularly lack power for rare variant association testing. We found that associations that are exclusively found by FuncRVP and not by the DeepRVAT burden test tended to be genes that were less tolerant to loss-of-function variants, on average (Fig. 5E).

### Novel associations identified by FuncRVP

FuncRVP identified 1,114 gene-trait associations (Table S3), of which 659 were shared with the DeepRVAT-based burden-test on the same training and validation samples and 455 were specific to FuncRVP (in contrast, DeepRVAT-based burden-test only had 166 specific associations). An additional 198 associations overlapped with findings from three large rare variant association studies on the UK Biobank, one comprising approximately 400,000 samples and the other two comprising about 470,000 samples^24,27,28^ (Fig. 6A), leaving 257 candidate novel associations. Among those, all but one agreed in direction on the test data, including 60 that showed nominally significant associations on the test samples (Fig. 6B, Table S4, adjusting for common variants located within 100 kb of the implicated genes, conditional association test, Methods). Moreover, most of these associations involved genes independently associated with common variants for the same traits (Fig. 6C, at least one lead SNP within 100 kb of the gene boundary^32^, 64.6% overlap vs. 21.9% genome-wide, *P* < 0.05). This evidence from independent analyses strongly supports the validity of the 257 novel associations identified by FuncRVP.

**Figure 6.**
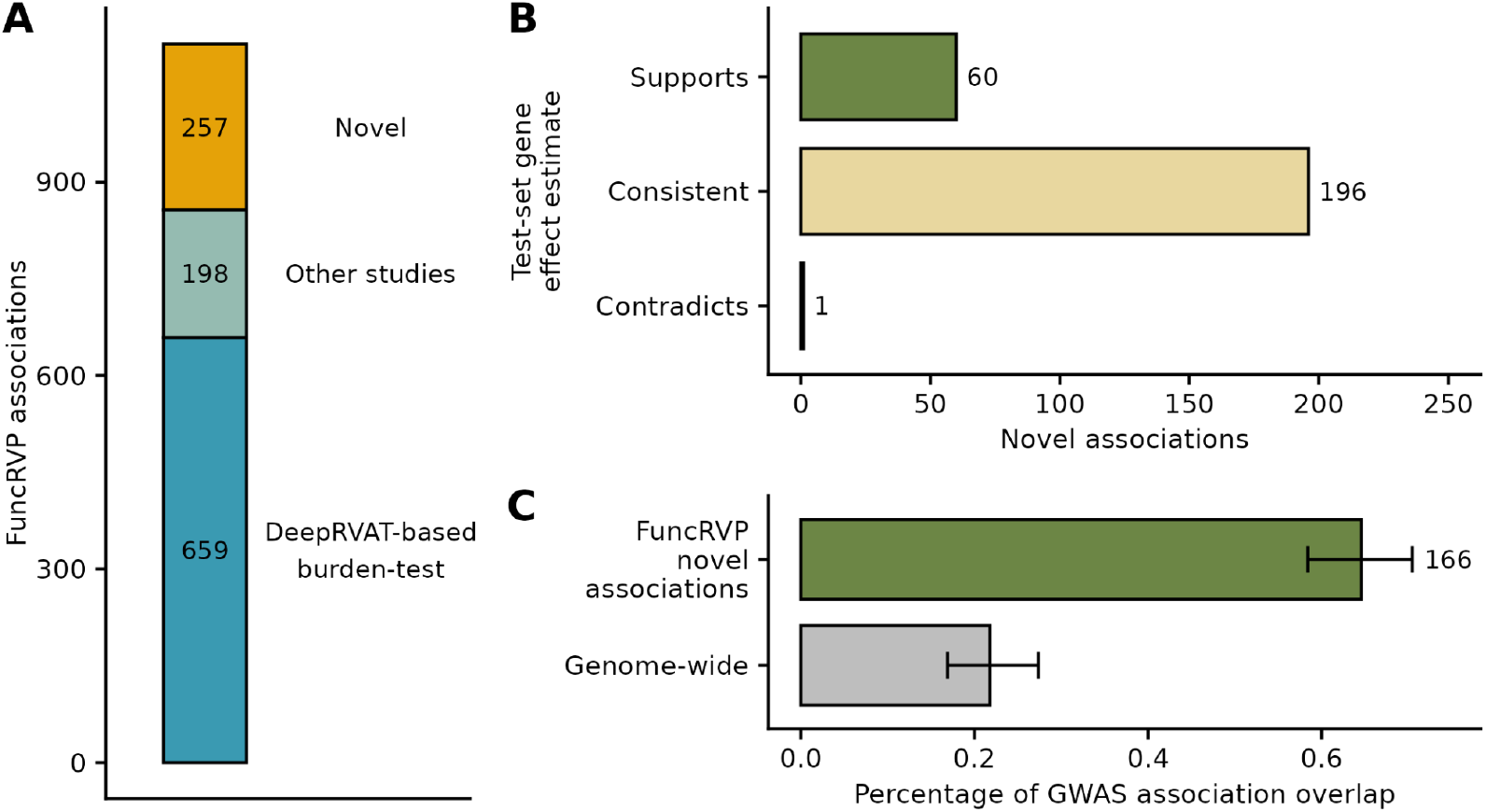
FuncRVP captures novel associations missed by other methods. **A)** The 1,114 associations of FuncRVP are split into three categories: if they are found by the DeepRVAT-based burden-test on the same data as FuncRVP, if they have been found by rare-variant association tests in previous studies (labeled here as “Other studies”)^24,27,28^, or if they are novel associations. **B)** Novel associations (y-axis) are categorized based on how they replicate on the test samples (x-axis). “Supports” if it is nominally significant on the test and in the same direction as the FuncRVP estimate. “Consistent” if the 95% confidence interval on the test samples overlaps the 95% credible interval of FuncRVP. “Contradicts” if the estimate on the test samples is nominally significant and in the opposite direction. **C)** Percentage of genes that have a significant GWAS variant in their vicinity (±100kb)^32^, computed for 257 FuncRVP novel associations and for genes chosen at random (“Genome-wide”). 166 out of 257 novel FuncRVP associations have GWAS overlap.

One notable finding was the negative association of impairment of *STAT5A* (Signal Transducer And Activator Of Transcription 5A) with mean corpuscular volume (MCV). In erythropoiesis, *STAT5A* is activated by erythropoietin signaling and is essential for red blood cell development^33^. Mouse knockout models of *STAT5A* exhibit significantly lower MCV^34–36^. This evidence, supported by held-out test samples and GWAS signals near the gene, strongly supports a causal role of impaired *STAT5A* on decreased mean corpuscular volume in humans (Fig. S2A).

Another interesting finding was the positive association between *SPTB* (Spectrin beta chain) impairment and high light scatter reticulocyte count. The dysfunction of *SPTB* can lead to hereditary spherocytosis^37^, a condition associated with significantly elevated reticulocyte counts^37,38^. This aligns with the positive effect found by FuncRVP, and that observed on the test data correcting for associated local common variants. The association is further strengthened by the importance of *SPTB* in providing mechanical support to the erythrocyte membrane skeleton^39^ (Fig. S2B).

We also confirm the positive association of impairment of *NPR1* and systolic blood pressure reported on the CHARGE+ Exome Chip BP cohort^40^, but not found on the UK biobank by previous studies (Fig. S2C).

## Discussion

FuncRVP advances rare variant association testing by leveraging gene impairment scores and functional gene information to enhance phenotype prediction and identify robust gene effect estimates on traits. Improved robustness and sensitivity are achieved by incorporating gene function information through embeddings. We found that the improvements could be achieved for a variety of embeddings, whereby those integrating gene co-expression appeared to perform best. Moreover, we found that the gene impairment score from DeepRVAT, which integrates multiple rare variant annotations, significantly outperformed the loss-of-function variant counts. Our analysis of 41 quantitative traits revealed 257 novel gene-trait associations with strong support from associations in held-out samples and enrichment for independent GWAS signal.

While FuncRVP outperforms the conventional linear regression-based methods, it is not without limitations. First, it models both the prior and the phenotype using a normal distribution, which can be restrictive when dealing with categorical traits, such as disease status, that cannot be accurately modeled with the normal distribution. Extending FuncRVP to handle categorical traits will require the use of approximate methods as conjugate priors for these distributions do not exist. Moreover, FuncRVP handles common variants associated with the trait in a simplistic way by including a common-variant polygenic risk score as a covariate. This approach does not leverage the functional relatedness between genes near trait-associated common variants and genes whose rare variant burden associates with the trait. Addressing this limitation in future work could enhance FuncRVP’s performance and complement emerging techniques integrating rare and common variants^41^. Another promising avenue for improvement is extending FuncRVP to jointly model multiple traits, allowing the method to leverage shared biological mechanisms between traits.

FuncRVP’s ability to exploit information from all genes was most striking when using the DeepRVAT impairment scores rather than the number of putative loss-of-function variants. The development of new impairment score models like DeepRVAT, AlphaMissense, and PrimateAI- 3D underscores the increasing availability of informative scores, making restrictions to the sole pLOF variants a less favorable option in most cases^6,13,42^. With the growth in the number and size of large-scale genome sequencing studies, along with improved rare variant annotations, FuncRVP’s ability to detect subtle genetic effects will be further enhanced, leading to the identification of a broader spectrum of genes contributing to human traits.

## Methods

### FuncRVP model

FuncRVP is a phenotype-specific model, with one model trained per trait. It predicts an individual’s trait measurement as a weighted sum of their rare variant gene impairment scores and covariates. More specifically, for an individual *i*, with gene impairment scores *g*_*i*_ (a vector of length equal to the number of protein-coding genes) and covariates *c*_*i*_, the trait measurement *y*_*i*_ is modeled as:

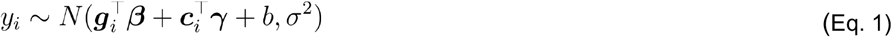

Where, *β* is the vector of all gene effects, *γ* is the vector of covariate effects, *b* is the intercept, and *σ*^2^ is the trait variance. We assumed a prior on the effect of all genes to be normally distributed, centered at zero, and with a gene-specific variance. Specifically, we assumed:

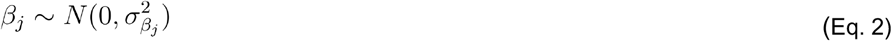

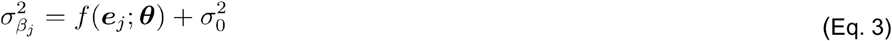

Where, *β*_*j*_ is the gene effect, 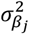 is the gene-specific prior variance, and the vector *e*_*j*_ is the functional embedding for gene *j*. The nonnegative function *f* is modeled using a neural network with parameters *θ*. The gene-specific variance 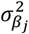 is given as the sum of a gene-specific term modeled by *f* on the embedding *e*_*j*_ and a constant minimal variance 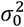 shared across all genes. To simplify notation, we use Z_*θ*_ to denote the diagonal matrix whose diagonal elements are the 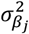 across all genes.

This Bayesian model was fitted in a two-step process. Initially, the parameterization of the neural network *f* was optimized by maximizing the likelihood of *y* (trait measurement vector for all samples) after marginalizing out the gene effects *β*, i.e.:

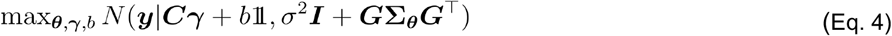

Where, ***G*** is the sample-by-gene matrix of all gene impairment scores, and ***C*** is the sample-by-covariate matrix of all covariates. We learn the optimal parameters 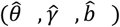 by minimizing the negative log-likelihood of *y* sub*j*ect to an L1-penalty on the parameters of *f*. We numerically minimized the resulting quantity using the Adam optimizer ^43^:

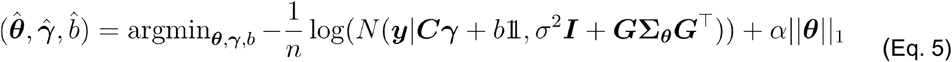

Where *n* is the number of samples in the batch. The learned gene effects after training were obtained from the posterior distribution of gene effects *β*|*y*, which is given by:

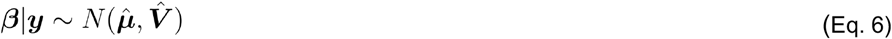

Where, the posterior mean 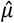 and posterior variance 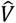. are given by:

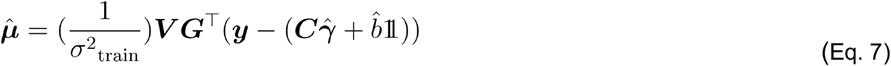

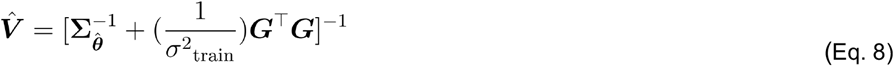

Where, 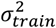 is the variance of the trait measurement observed on the training samples, and 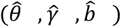are the learned parameter values on the training data. To compute the posterior distribution of gene effects, we used trait measurements *y*, genotypes ***G***, and covariates ***C*** from the training and validation samples. The gene effect posterior mean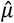 thus obtained was reported as the effect of the gene on the trait. The gene effect posterior mean 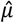 and variance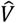. were then fixed and used to obtain phenotype predictions on the held-out test data.

Given a new, unseen sample with genotype *g*^*ne*w^, and covariates *c*^*ne*w^, the phenotype *y*^*ne*w^ was predicted to distribute according to:

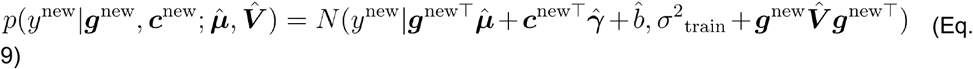

We used the expected value of *y*^*ne*w^ from (Eq. 9) as FuncRVP’s prediction of the trait measurement for the new sample.

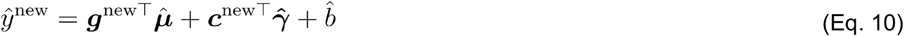

### Covariates

We considered for each trait age, age^2^, sex, age x sex, age^2^ x sex, the first 20 genetic principal components, and the common variant polygenic risk score as the covariates in the model.

### Train-validation-test split

The 372,392 samples from the UK Biobank were divided into train, validation, and test sets in a 65:10:25 ratio, respectively. The train set was utilized to learn the model parameters, the validation set was employed to optimize the hyperparameters, and the test set was used to evaluate the phenotype prediction performance of the model. The same train, validation, and test split was consistently applied across all models.

For the initial assessment of model assumptions, i.e., the comparison between modeling the expectation of the prior against the variance (Fig. 2A), a smaller dataset of 161,850 with a train, validation, and test sets in an 80:20:20 ratio was used.

### Parameter optimization

The neural network *f* consisted of a variable number of layers. Softplus activation functions were applied between all layers to introduce non-linearity and after the last layer to ensure positive output values. The constant minimal variance 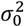 was initialized at 10^−5^. We optimized the model using the Adam optimizer ^43^, with a learning rate of 10^−3^. Training was performed for 50 epochs, with a batch size of 16,384 samples. The model performance was monitored on a validation dataset and the parameters at the epoch with the highest validation R^2^ were saved.

### Hyperparameter optimization

The best model architecture for each trait was determined through hyperparameter optimization. Five trials were conducted for each combination of hyperparameters, and the second lowest validation R^2^ was selected as a metric for the performance of that particular combination. We found that selecting the second lowest validation R^2^ led to more robust results than selecting the highest R^2^. Hyperparameter tuning was accomplished using a TPE (Tree-structured Parzen Estimator) algorithm ^44^, implemented via the Optuna Python package ^45^, to optimize the number of hidden layers : [1, 4], size of hidden layers : [2,12], L_1_ regularization strength, ***α*** : [10^−10^, 10], activation function used: {Softplus, ReLU}, the initialization of the bias term of the last hidden layer : [-10,10], and the initialization of the constant minimal variance, 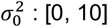.

### Last-layer bias

We found the bias of the last layer of the neural network *f* to be important for model convergence. Hence, we included the initialization of this parameter as a hyperparameter. A larger last-layer bias would allow more genes to have non-zero prior variances and, therefore, a higher variance of the prior.

### Posterior of gene effects

To rank gene associations obtained by FuncRVP, we compute the probability of direction^46^ on the posterior distribution of all gene effects. The probability of direction indicates the certainty in the sign of an effect and lies between 0.5 and 1. Given the posterior distribution of gene effects *p*_*post*_ on a variable *β*, the probability of direction *p*_*d*_ is given by:

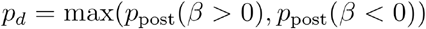

### Baseline models

#### Common-variant baseline model

To evaluate the improvement in phenotype prediction due to rare variants, we consider a baseline model capturing the effects of covariates and common variants through the polygenic risk score. The common-variant baseline phenotype predictor is given by:

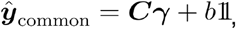

where ***C*** is the matrix of covariates including the PRS and *b* is the intercept. Since this model requires no hyperparameter optimization, it is trained on all training-phase samples, i.e., the union of the train and validation splits of FuncRVP.

#### Burden test

Burden tests aggregate variant effects in a region, usually a gene. These aggregated effects are known as the gene impairment scores (or gene burdens), and these are used to test the association of the gene with a trait^10^. For a given trait measurement *y* and gene *j*, we perform the burden test using a linear regression with the covariates:

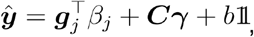

where *g*_**j**_ is the DeepRVAT gene impairment score of gene *j* for all individuals, ***C*** is the matrix of covariates including the PRS and *b* is the intercept. Since this model requires no hyperparameter optimization, it is trained on all training-phase samples, i.e., the union of the train and validation splits of FuncRVP.

#### Phenotype prediction linear model

After running the burden test across all genes, we selected the genes significantly associated with the phenotype using Bonferroni significant p-values at 0.05 (multiple testing correction performed per trait). We then used only these significant genes to train a linear model for phenotype prediction:

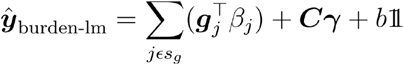

where *s*_*g*_ is the set of significant genes for the phenotype. Since this model requires no hyperparameter optimization, it was trained on all training phase samples, i.e., the union of the train and validation splits of FuncRVP.

#### FuncRVP, randomized embedding

We used a randomly generated embedding as a negative control experiment. Specifcially, funcRVP was trained in the same manner as described above, with the exception that the embeddings were generated through isotropic Gaussian distributions. The training and evaluation procedures were identical to FuncRVP.

### Model evaluation

#### Comparison of phenotype prediction

FuncRVP predicts the phenotype given an individual’s genotype. We trained independent models for each phenotype and evaluated them on a held-out test set by comparing phenotype prediction R^2^ against the baseline models. Predictions on the test set were computed using the posterior of the gene effects obtained after the training procedure. We used the relative improvement in explained variance compared to the common-variant baseline model as our metric. This allowed us to compare the performance increments due to rare variants. Given the explained variance of a model on the held-out test data (*R*^2^ _model_) and that of the common-variant baseline model (*R*^2^ _common_), the final metric Relative *ΔR*^2^ was defined as:

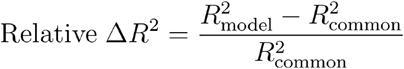

We conducted a bootstrapping to identify traits where the two models perform significantly differently. Specifically, we drew 1,000 bootstraps from the test set and computed the phenotype prediction R^2^. We computed the p-value of these comparisons as follows ^47^:

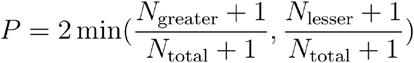

where *N*_greater_ is the number of bootstraps where FuncRVP performed better, *N*_lesser_ is the number of bootstraps where FuncRVP performed worse, and *N*_total_ is the total number of bootstraps drawn, which was always 1,000. We consider *P* < 0.05 as the significance threshold.

#### Conditional association tests

Conditional association testing was performed to control for possible confounding due to common variants in linkage with the FuncRVP-discovered genes (candidate gene-trait association, probability of direction >0.999). To this end, independently associated common variants from Pan-UKBB^32^ GWAS summary statistics were identified through LD-based clumping using PLINK^48^ (v1.9) with default parameters, restricting to associations with a p-value less than 10^™7^ and a minor allele frequency larger than 1%. Next, each candidate gene-trait association was statistically assessed on the test data using linear regression of the trait against the gene impairment score, the independently associated common variants, the PRS, and the covariates. Nominally significant associations (F-test, *P* < 0.05) were retained for further investigations.

#### Novel associations test set support

We compared gene effect estimates of the novel associations of FuncRVP obtained on the training and validation split samples to the gene effect estimates obtained by a burden test on the test split samples. We categorized them into three categories (Supports, Consistent, and Contradicts) based on the strength of the overlap as follows. An effect was supported of the test- set estimate was nominally significant on the test samples and in the same direction as the FuncRVP estimate; It was considered to be consistent if the 95% confidence interval on the test samples overlapped the 95% credible interval of FuncRVP; and it was contradicted if the test-set estimate was nominally significant but in the opposite direction of the FuncRVP estimate.

### Datasets

#### Phenotype processing and filtering samples

We used 469,382 whole-exome sequencing samples with corresponding blood and urine measurements from the UK Biobank^26^ for training and benchmarking FuncRVP phenotype prediction performance against other approaches. We used the *ukb_gen_samples_to_remove* function of the ukbtools R-package (v0.11.3^49^) together with pre-computed relatedness scores (see UK Biobank Resource 668) to remove closely related individuals, keeping only one representative of groups that are related to the 3^rd^ degree or less. Individuals with European ancestry were identified using UK Biobank data field 22006 (termed ‘Caucasian’). This filtering resulted in a dataset of 372,392 individuals.

All measurements were inverse-rank-normal-transformed, corresponding to the data preprocessing in the Genebass study^27^. For each trait, the dataset of genotypes, covariates, and corresponding trait measurements was split randomly into training, validation, and test data containing 65%, 10%, and 25% of the data, respectively.

#### Covariates and Polygenic Risk Scores (PRS)

For each sample, the covariates: age (UKBB data field 21003), age^2^, sex (UKBB data field 31), age x sex, age^2^ x sex, and the first 20 genetic principal components (UKBB data field 22009_0.[1- 20]) were obtained directly from the UK Biobank. All of these were used as covariates for the burden test and for FuncRVP. Polygenic risk scores were obtained from the PGS catalog^50^ using the study by Privé et al.^51^ and applied to the genotyping array data using PLINK 2.0^52^.

#### Gene impairment scores

We primarily used the DeepRVAT gene impairment scores for all models^13^. DeepRVAT summarizes multiple variant effect annotations of all variants considered rare (allele frequency, AF, less than 0.1%) into one aggregated score per gene. The DeepRVAT gene impairment scores were obtained for 469,382 individuals from the UK Biobank for 17,984 protein-coding genes as described earlier^24^. DeepRVAT gene impairment scores were median shifted to center the median score of each gene to zero. This ensured that a zero gene impairment resulted in no effect of the gene on the phenotype and was consistent with the regularizing prior.

Alternatively, we considered as gene impairment score the number of putative loss-of-function (pLOF) variants, defined as any variant having the following consequences by Ensembl VEP^25^ on the canonical transcript: stop gained, start lost, splice donor, splice acceptor, stop lost, or frameshift.

#### Gene Embedding

We utilized gene function representations from Brechtmann et al.^20^. In these functional gene embeddings, each protein-coding gene was encoded as a 256-dimension real vector. The PoPS feature matrix had 57,743 dimensions, which were projected to 256 dimensions using PCA. The concatenated “Omics + PoPS” gene embedding had 512 dimensions, 256 dimensions from Omics and 256 from PoPS. Upon intersection with the genotype matrix from DeepRVAT, we obtained 17,517 coding genes.

The “Enformer” embedding was derived from Enformer^30^, a sequence-based model predicting gene expression and chromatin states across tens of cell lines and conditions. Embeddings were extracted at each gene’s Ensembl canonical transcription start site (TSS), resulting in 3072- dimensional vectors. We reduce the dimensionality of this embedding by using the first 512 principal components, which captured 95.53% of the variance.

The “ESM2” embeddings were obtained from the protein language model ESM2^29^. For each gene, we extracted the 1280-dimensional latent representation of the model’s final layer. We reduce the dimensionality of this embedding by using the first 512 principal components, which captured 98.65% of the variance.

The “gene2vec” embedding was obtained from the original publication^21^. It is an embedding of gene co-expression made from 984 whole transcriptome human gene expression data sets from Gene Expression Omnibus (GEO)^53^. This embedding has a 200-dimensional representation of each gene, requiring no further dimensionality reduction.

#### Discoveries from other rare-variant association studies

We included gene-trait associations found by the genebass study^27^ and the study by Backman et al.^28^. These rare-variant association studies were performed on 394,841 and 454,787 individuals from the UK Biobank. Gene-trait associations for all 41 traits that we used were available in both studies.

#### Overlap of discovered genes with GWAS results

On the Pan-UK Biobank^32^ GWAS summary statistics, we identified independently associated variants through LD-based clumping using PLINK v1.9^48^ with default parameters. These variants were filtered at the conventional GWAS nominal p-value threshold of 5 × 10^−8^ to obtain the index variants for each phenotype.

The genes with an index variant within ±100kb of the gene were identified for each trait. We computed *n*_trait_, which is the number of these genes which overlap with FuncRVP gene-trait associations. An empirical p-value was obtained for each trait by drawing 10,000 bootstraps with replacement of *n*_trait_ genes from the set of all protein-coding genes and computing the overlap with the set of genes with a GWAS index variant in their vicinity.

#### Burden heritability

Burden heritability scores were obtained from supplementary table 8 of the work by Weiner et al.^2^. The column “aggregated_h2” from this table was used as the burden heritability score for each trait. The traits studied here overlapped with only 16 out of the 41 traits that we benchmark on.

#### LOEUF scores

The loss-of-function observed/expected upper bound fraction (LOEUF) was obtained from the download portal of gnomad v2.1.1^1^.

## Computing resources

The FuncRVP model trainings were capped at 50 epochs as more epochs led to minimal performance improvements on the validation set. The entire training procedure took typically a little more than 30 minutes on one NVIDIA A40 for each trait.

## Acknowledgments

We thank Brian Clarke, Marcel Mück, and Magnus Wahlberg for the computation of DeepRVAT gene impairment scores. We thank Johannes Hingerl for extracting gene embeddings from Enformer and ESM2. S.L. thanks Vicente Yépez for feedback and advice.

This research has been conducted using data from UK Biobank, a major biomedical database (project IDs 25214 and 81358). This study was funded by the Deutsche Forschungsgemeinschaft (DFG, German Research Foundation) via the project NFDI 1/1 “GHGA - German Human Genome-Phenome Archive” (#441914366 to S.L., E.H. and J.G.), by the German Bundesministerium für Bildung und Forschung (BMBF) through the ERA PerMed project PerMiM (01KU2016B to S.L., J.L., F.B., and J.G.) and through the Model Exchange for Regulatory Genomics project MERGE (031L0174A to F.R.H. and J.G.). This study was supported by the Deutsche Forschungsgemeinschaft (DFG, German Research Foundation) via the IT Infrastructure for Computational Molecular Medicine (project #461264291). This work was funded by the Deutsche Forschungsgemeinschaft (DFG, German Research Foundation) through the TRR267 (403584255, sub-projects, Z03 to J.G. and B05 to Z.C.). This work was funded via the EVUK programme (“Next-generation Al for Integrated Diagnostics”) of the Free State of Bavaria (to F.B. and J.G.). This work was funded by the Initiative and Networking Fund of the Helmholtz Association in the framework of the Helmholtz AI project call (project DeepVar, ZT-I-PF-5-156). S.L. and E.H. are supported by the Helmholtz Association under the joint research school ‘Munich School for Data Science – MUDS’. F.P.C. was funded by the Free State of Bavaria’s Hightech Agenda through the Institute of AI for Health (AIH). The Ethics Committee of the Technical University of Munich issued a statement of no objection regarding this study (reference #2025- 263-S-KK).

## Contributions

F.B. and J.G. conceptualized the project. F.B., F.P.C., and J.G. designed the methodology. J.L. and S.L. implemented the methods. S.L., J.G., and F.B. analyzed the data and created the visualizations. F.R.H., E.H., J.L., and Z.C. curated the data. S.L. and J.G. wrote the original draft of the manuscript. J.G. supervised the project.

## Corresponding author

Correspondence to Julien Gagneur and Felix Brechtmann.

## Data and code availability

The code is made available on GitHub: https://github.com/gagneurlab/funcRVP

