## Supplementary Figures for "Functional gene embeddings improve rare variant polygenic risk scores"

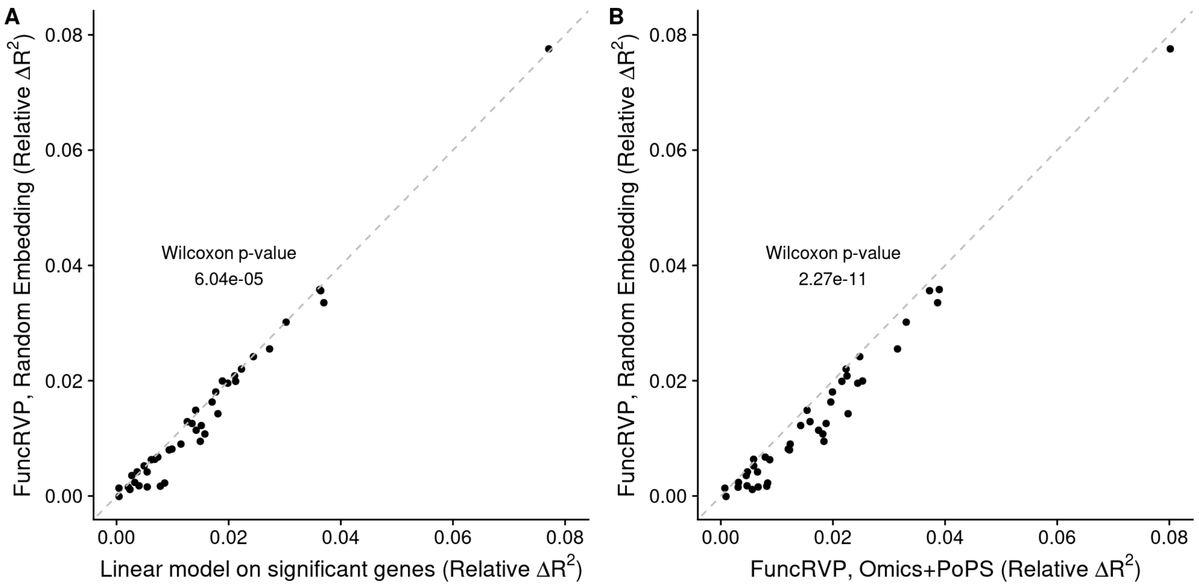


**Figure S1. FuncRVP with a randomly generated embedding performs worse than linear regression on burden-test significant genes. A)** Explained variance on held-out data using FuncRVP using a randomly generated embedding (y-axis) compared to a linear regression on gene impairment scores of significant genes (x-axis) for 41 phenotypes. The relative fit improvement was computed as the improvement in $R^2$ relative to a baseline linear model using only the PRS and covariates. Overall significance was determined through a paired Wilcoxon test. **B)** Same as in (**A**), except FuncRVP using a randomly generated embedding (y-axis) is compared against FuncRVP with the Omics+PoPS embedding (x-axis).

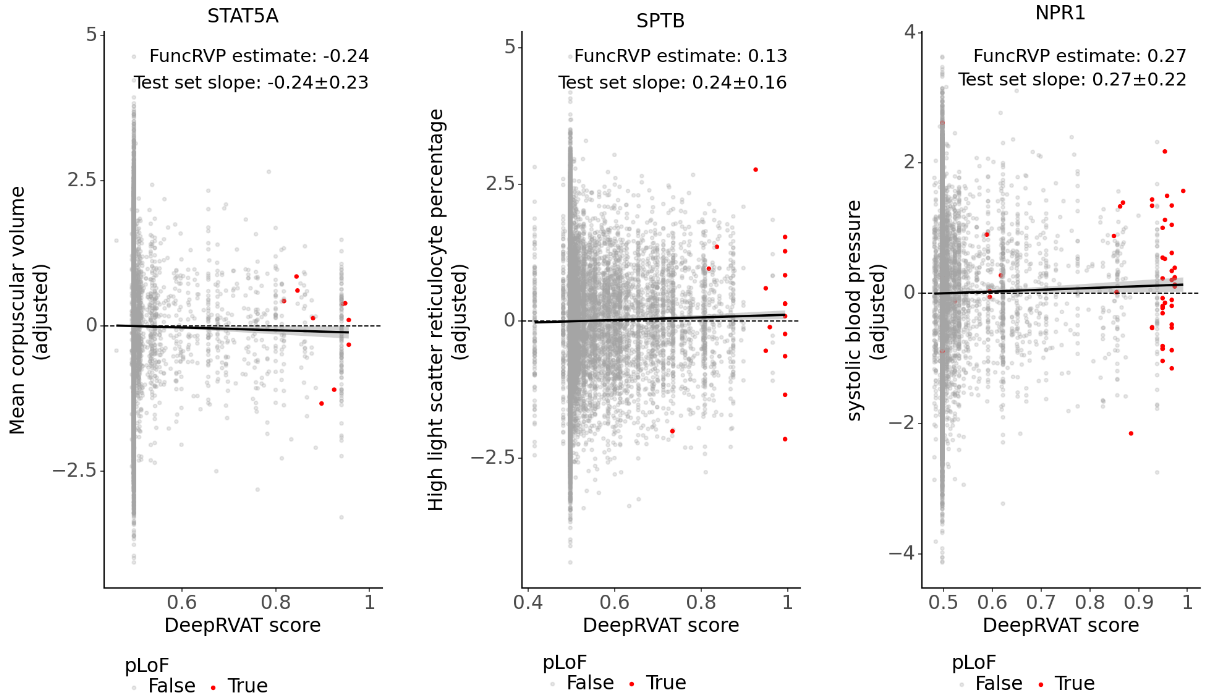


**Figure S2. A)** Mean corpuscular volume adjusted for local GWAS signal, PRS, and covariates (y-axis) plotted against the *STAT5A* DeepRVAT gene impairment score (x-axis) for all individuals in the test samples for the three novel associations described in the text. Individuals with pLoF variants in the gene are highlighted in red. The trend line is plotted using linear regression on the test samples. The “FuncRVP estimate” indicates the gene effect estimated by FuncRVP, and the “Test set slope” indicates the slope of the linear regression with a 95% confidence interval computed on the test samples. **B)** Same as in (**A**), but for the association between the phenotype High light scatter reticulocyte count and impairment of the gene *SPTB*. **C)** Same as in (**A**), but for the association between the phenotype systolic blood pressure and impairment of the gene *NPR1*.
